# Hyper-connectivity of the striatum related to restricted and repetitive behaviors’ severity in children with ASD

**DOI:** 10.1101/2020.02.21.957993

**Authors:** Irene Dupong, Adriana Di Martino

## Abstract

Restricted and repetitive behaviors (RRB) are a core feature of autism spectrum disorder (ASD). But little is known about the underlying neurobiology of the disorder, preventing from having specific therapeutic targets. Based on the literature, we explored the correlates between a clinical score of RRB, using the Repetitive Behaviour Scale – Revised, and the intrinsic connectivity of seven striatal regions in a sample of 157 children with ASD. The sample was acquired from the ABIDE consortium. We found a significant correlation between the severity of our clinical scale and several cortico-striatal networks. Specifically, stronger connections were found between striatal seeds and two cortical areas, an occipital area and a frontal area in the left hemisphere. Intrinsic functional connectivity of the striatum could serve as a potential biomarker for improved detection of RRB severity.

## Introduction

Restricted and repetitive behaviors (RRB) are a core feature of autism spectrum disorders (ASD). They include : stereotypies, insistence on sameness, inflexible routines, or ritualized patterns or verbal nonverbal behavior, highly restricted, fixated interests and hyper- or hypo-reactivity to sensory input or unusual interests in sensory aspects (1).

Despite an emerging number of studies on the subject, the underlying neural basis for these features remain incompletely understood. Few studies have focused on RRB, authors favoring symptoms of social impairments, the other core feature of ASD (1). To asses RRB a frequently used scale is the Repetitive Behavior Scale-Revised (RBS-R), that captures symptom scope at different ages (2).

Theoretical understanding and neuroimaging studies have associated RRB and ASD with the striatum, a subcortical structure belonging to the basal ganglia associated to a variety of functions including mouvement, learning, cognition and emotion (3–5). More precisely, the neuroimaging studies have associated RRB to striatal volume and connectivity anomalies in patients with ASD compared to typical developing controls (6–15). Without taking into account RRB, the striatum has been associated with ASD in functional connectivity studies (16,17). Thus, investigating the connectivity between the striatum and other brain regions may be key to understanding the nature of restricted and repetitive behaviors in ASD.

We used resting state functional connectivity magnetic resonance imaging (MRI) to examine the intrinsic functional connectivity of striatum in an international cohort of children and adolescents with ASD. Quantifying the synchronization of blood oxygen level dependent (BOLD) signal fluctuations between different brain regions, has detected the functional networks that echo with many known features of anatomical organization in subjects with ASD, in other disorders and in healthy subjects (18–21). More precisely, studies using functional MRI (fMRI) in ASD have reported numerous abnormalities in intrinsic functional connectivity (iFC) as compared to those in typically developing controls, ranging from hypo-connectivity (22–24) to hyper-connectivity (16,25–29) or both (30,31). These inconsistencies are attributed to the heterogeneity of the cohorts, small sample sizes, methodological variations and heterogeneous diagnostic methods.

The present study investigated iFC of the striatum, in children and adolescents with ASD, by using resting-state fMRI. We hypothesized that children with ASD would exhibit striatal connectivity patterns that correlate with clinical scores of their RRB. A further aim was to localise the areas, and correlate them with other clinical features.

## Material and Methods

### Sample selection

The current sample was selected from the ABIDE II repository (32). Consistent with the primary aim of this study, we included only MRI data from individuals with a diagnosis of ASD and available RBS-R scores that met the following criteria: 1) RBS-R completed by an informant as differences in self and informant ratings may confound results (2) ; 2) mean frame-wise displacement (mFD) < 0.6 mm as this was 2 standard deviations (SD) from the mean of the originally selected sample (0.172 mm ± 0.21 mm) (33); 3) successful functional to anatomical registration. After taking the above steps, we confirmed that each collection presented with at least four datasets meeting these criterias. These steps yielded a final ASD sample of 157 datasets. Data selection flow and sample characteristics are detailed in Supplementary Figure 1 and Table 1, respectively.

**Table 1:**
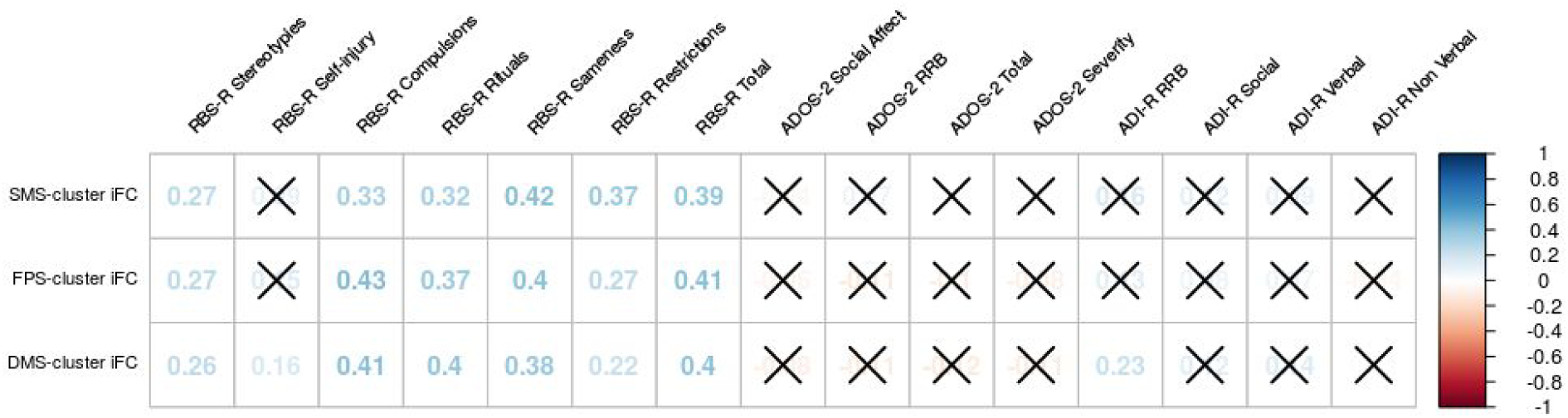
Correlations between residual z-scores of the iFC of each seed-cluster group and each score of the RBS-R scale as well as total and measures of ADOS-2 and ADI-R. Crossed numbers are correlations with a p value > 0.05.

### Imaging acquisition

Given that the ABIDE II cohort results from a multi-site retrospective data aggregation, sequence parameters for anatomical and functional MRI data varied. Though all data were collected with 3 Tesla scanners. Details regarding data acquisition for each sample have been provided in the ABIDE website [http://fcon_1000.projects.nitrc.org/indi/abide] and are summarized elsewhere (32).

### Preprocessing

Data were analyzed using version 0.3.9.1 of the Configurable Pipeline for the Analysis of Connectomes (C-PAC, http://fcp-indi.github.com/C-PAC) (34), which integrates tools from AFNI (http://afni.nimh.nih.gov/afni), FSL (http://fmrib.ox.ac.uk) and Advanced Normalization Tools (ANTs; http://stnava.github.io/ANTs) using Nipype (http://nipype.readthedocs.io/en/latest/). Before skull-stripping (using FSL BET), all anatomical images were spatially normalized to MNI152 2mm stereotactic space with linear and non-linear registrations using ANTs (35). For all functional images we performed site-specific slice time correction (AFNI 3dTshift commands) based on each site acquisition parameters. Motion correction was performed using the AFNI command 3dvolreg by a two-pass procedure. In a first step, each functional volume was co-registered to the (un-aligned) mean functional image. In a second step, a new functional mean image based on the aligned images was used as the reference image. At this second stage, motion parameters based on the Friston 24-Parameter Model (six motion parameters, their values of preceding volumes, 12 squared values of these items (36)) were regressed. Nuisance regression also included the mean WM and CSF signals obtained in subject-specific masks with tissue probability thresholds of 0.96 and linear and quadratic trends. Following nuisance regression a band-pass filtering (0.01-0.1 Hz) was applied using the AFNI command 3dBandpass. A functional-to-anatomical co-registration was achieved by Boundary Based Registration (BBR) using FSL FLIRT (37). Spatial normalization of functional EPIs to MNI152 space was done by applying linear and non-linear transforms from ANTs.

### Seed definition and analyses

Given that the striatum’s regions have heterogeneous functions, we based our seed given the striatum’s iFC. At the individual level, we measured the iFC of different striatal seeds, chosen a priori as regions-of-interest. We based our masks on the seven striatal functional divisions empirically validated by Choi and colleagues (38). We selected these functional divisions over others as each of them map to known cortical networks including the visual, somato-motor, dorsal and ventral attention, limbic, fronto-parietal and default mode network (39). We included all these striatal divisions as each of the striato-cortical network they belong have been implicated in the processes affected by ASD (25,40–43). The average time series of all voxels within each ROI was individually computed, and correlations between these time series and those of all other brain voxels were assessed. Whole-brain voxels were defined by a study-specific volume masks including voxels in MNI space present across 99% of all individual data. The resultant connectivity maps were smoothed (applied full-width at half-maximum 4 mm). This resulted in individual participant-level maps of all voxels exhibiting significant iFC with each of the seven seeds.

### Group-level analysis

A general linear model was fitted at each voxel that included five nuisance covariates (site, sex, demeaned age, demeanded mFD (33), and demeaned whole brain iFC values) and RBS-R Total scores as the primary covariate of interest. This analysis generated maps of regions exhibiting significant positive and negative FC for each seed. Gaussian random field theory was used for cluster-level multiple comparisons correction (Z ≥ 2.58; p ≤ 0.05). In secondary analyses, group analyses were repeated using a more stringent threshold of Z > 3.1.

### Post-hoc analysis

To localise the clusters, maps were used in FSL to extract the percentage of voxel in the Talairach structural atlas and in Yeo’s functional atlas.

Correlation analyses were applied to examine whether iFC within clusters was associated IQ, subscores of RBS-R, medication status (44), comorbidities and other clinical (ADOS-2 and ADI-R) indices, when data was available data. Finally, we divided the ASD group into low and high RBS-R total scores using the mean of the sample (n = 21) as the threshold. We also extracted iFC for the sames striatal seeds on a TDC population of 201 subjects, with the same inclusion criteria, image acquisition, preprocessing and seed definition analysis.

## Results

Our sample comprises 157 subjects ASD (mean age of 10,77 years, 76% males), supplementary characteristics are shown in Supplementary Table 1.

### RBS-R striatal iFC relationships

Analyses revealed significant associations between the RBS-R scores and the coefficient of IFC between clusters and three striatal seeds. Specifically, as shown in Figure 1, RBS-R total scores were positively associated with the iFC between the striatal somato-motor seed and a cluster in the cuneal cortex (occipital cortex) on both hemispheres. Positive relationships were also observed for the iFC of the fronto-parietal and default mode striatal seeds with partially overlapping regions of the superior frontal gyrus in the left hemisphere (see Supplementary Table 2).

**Figure 1:**
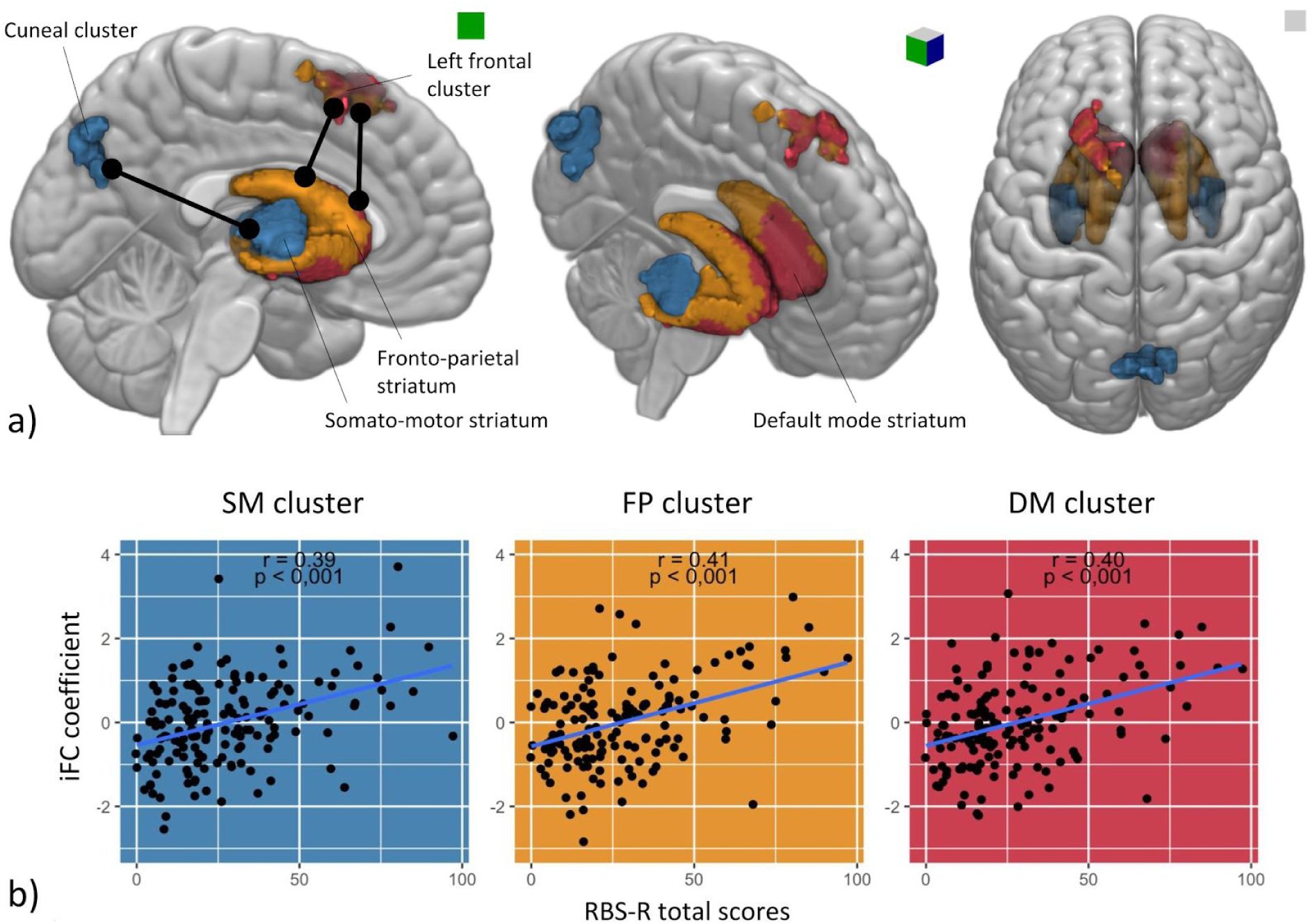
Correlations between significant intrinsic functional connectivity clusters (z values) and RBS-R total scores. a) Anatomical localization of the associations b) Scatterplots : r values represent correlations between z scores for each of the three clusters of significance and the RBS-R total scores for each subject. Significant results were obtained for the same criteria with Z = 3.1, but for a smaller cluster.

### Functional localization of the clusters

Following Yeo’s functional cortical atlas (39), the cluster associated with the SM striatal seed is distributed amongst several functional cortical areas, including mainly the visual cortex, the dorsal attention cortex, de fronto-parietal cortex and overlapping on some white matter. Whereas, the clusters associated with the FP and DM striatal seeds belong mostly to the default mode cortical area and to a lesser extent to the fronto-parietal areas. The distribution is shown in Figure 2.

**Figure 2:**
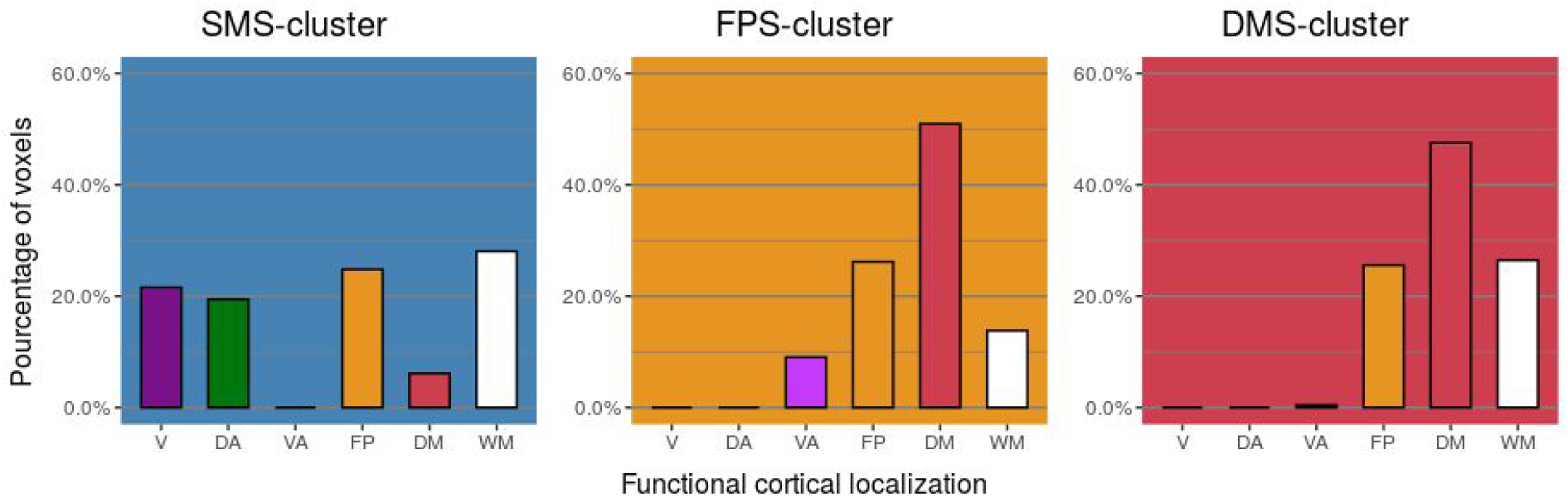
Distribution of each of the clusters’ voxels amongst Yeo’s atlas. DA : dorsal attention cortex, DM: default mode cortex, DM: default mode striatal seed, FP : fronto-parietal cortex, FPS : fronto-parietal striatal seed, SMS : somato-motor striatal seed, V : visual cortex, VA : ventral attention cortex.

### Correlations with clinical features

Correlational analyses were used to examine whether connectivity between the striatal seeds and the clusters showing significant differences in regard to RBS-R total scores were influenced by intelligent quotient (IQ), comorbidity or medications.

In our sample IQ was correlated to RBS-R total scores. Global IQ was not available for all individual and was not included in the initial analysis. Is subjects with available global IQ, we used the same general linear model integrating the z-scores for the 3 strial seed and cluster groups, the five nuisance covariates and global IQ scores. We also integrated in another calculation the Performance IQ, which was available for all individuals. Then, Pearson correlations were generated between the residual iFC of each group and RBS-R total scores. As shown in Figure 3, correlation are still significant even when integrating IQ scores.

**Figure 3:**
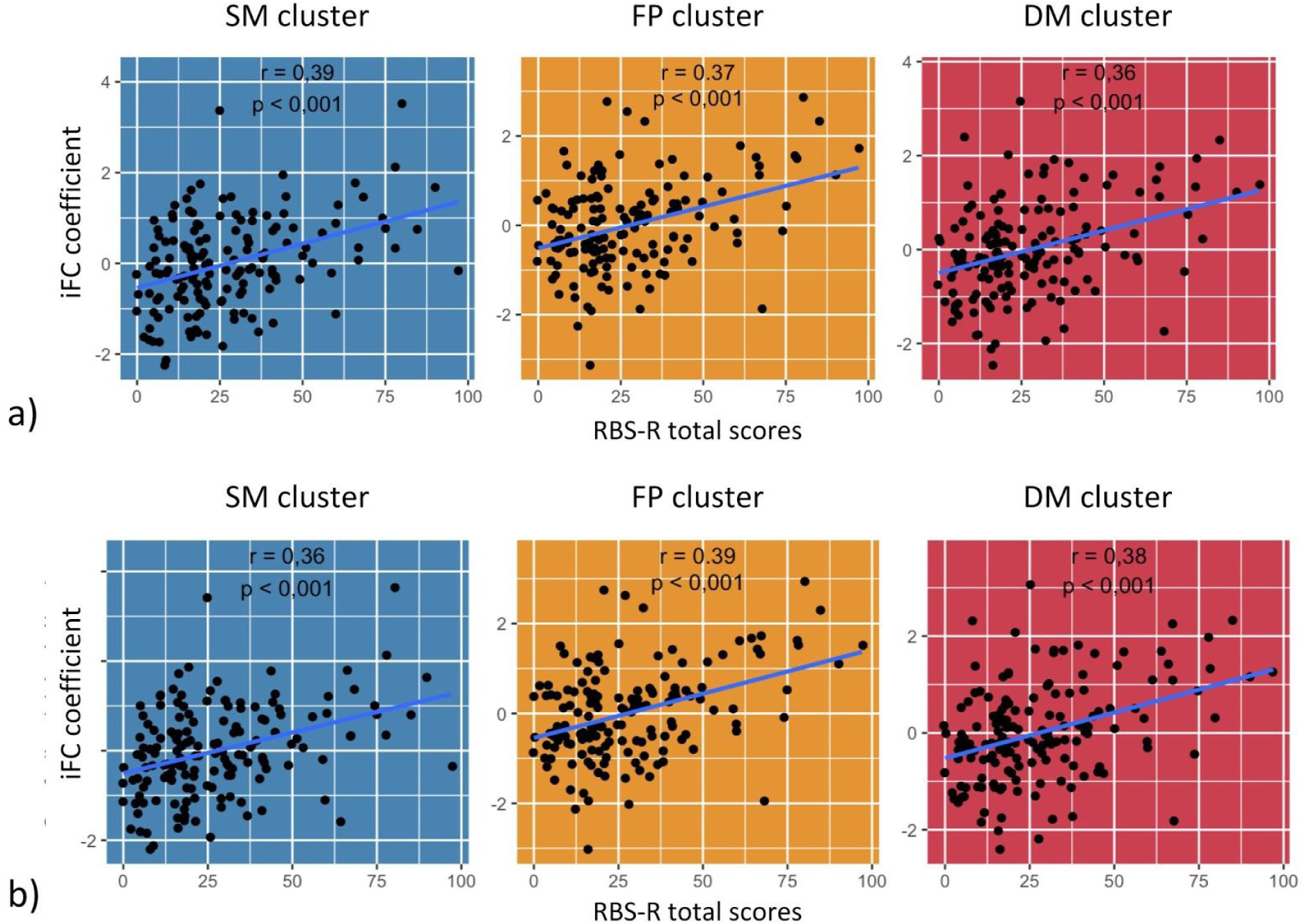
Scatterplots of correlation between residual z-scores of the iFC of each seed-cluster group and RBS-R total scores, when including a) Global IQ and b) Performance IQ.

To assess if medication use and/or comorbidities had an impact on our results, we used the same model previously used but instead of including IQ we included the medication and/or comorbidity status. As shown in figure 4, RBS-R total scores are still significantly correlated with the iFC of each seed-cluster when considering medication and/or comorbidity, using Pearson correlations. Only 101 subjects out of the 157 had information regarding comorbidities.

**Figure 4:**
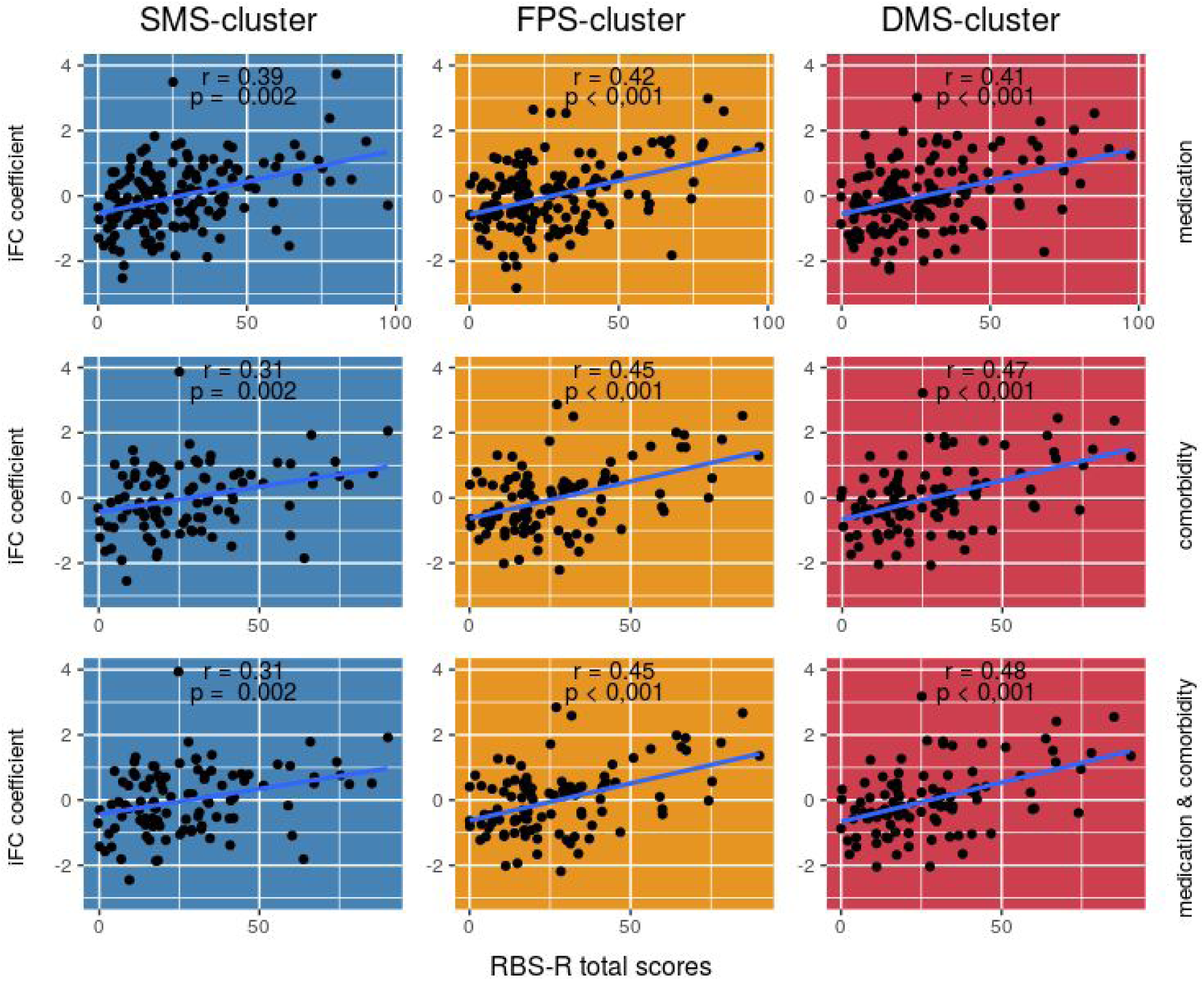
Scatterplots of correlation between residual z-scores of the iFC of each seed-cluster group and RBS-R total scores, when including a) medication use, b) comorbidities and c) medication use and comorbidities.

### Correlations with other RRB scores and measures of autism symptoms (RBS-R subscores, ADOS and ADI-R)

First, since RBS-R total scores regroup a heterogenous set of symptoms, assessing how RBS-R subscales weighted on the correlation between RBS-R total and the iFC seemed important. Pearson correlations were generated between the residual iFC (integrating z-scores and the five nuisances) of each seed-cluster and each of the 6 RBS-R subscales (stereotyped behavior, self-injurious behavior, compulsive behavior, ritualistic behavior, insistence on sameness and restricted behavior). Five of the six subscales correlated with all the seed-cluster iFC. The subscale for self-injurious behavior was only correlated with the iFC between the DMS seed and it’s cluster. The iFC between the SMS seed and it’s related cluster is mainly positively correlated to sameness. Whereas the one between the FPS and DMS seeds are most correlated with compulsive behaviors. Secondly, we were interested on how the iFC was related to diagnostic assessment scales of ASD. Out of our 157 subjects, 122 subjects had total and subscores available for ADOS-2 and 127 subjects for ADI total and subscores. Again, Pearson correlations were generated between the residual iFC (integrating z-scores and the five nuisances) of each seed-cluster and domains for ADOS-2 (Social Affect subtotal, Restricted and Repetitive Behaviors subtotal, Severity subtotal and Total score) and ADI-R (Restricted and Repetitive Behaviors subtotal, Reciprocal Social Interaction subtotal, Verbal Communication subtotal and Non Verbal Communication subtotal). Only the activity of the DM striatal seed and it’s cluster were corelated to ADI-R RRB scores. In our sample, RBS-R total scores were correlated with ADI-R RRB scores. Table 1 shows correlations between all RBS-R scores, the other ASD clinical scores and the seed-cluster iFC.

### High RBS-R ASD subject vs Low RBS-R ASD subject vs TDC

Finally, we divided the ASD group into low-RBS-R total scores and high-RBS-R total scores and compared there results amongst them and with the sample of TDC. Regarding the TDC sample, there were no group differences on age or handedness or eye status during the scan (opened or closed). But the samples did differ on IQ scores, as expected on RBS-R total scores and medication status. There were more males in the ASD group.

With each of these three groups, Pearson correlations were generated between the residual iFC (integrating z-scores and the five nuisances) of each seed-cluster and RBS-R total scores. As shown in Figure 5, only the High RBS-R ASD sample was significantly correlated with the all three iFC for cluster-seed groups. The low RBS-R ASD sample was only correlated with the iFC of the SM striatal seed and cluster.

**Figure 5:**
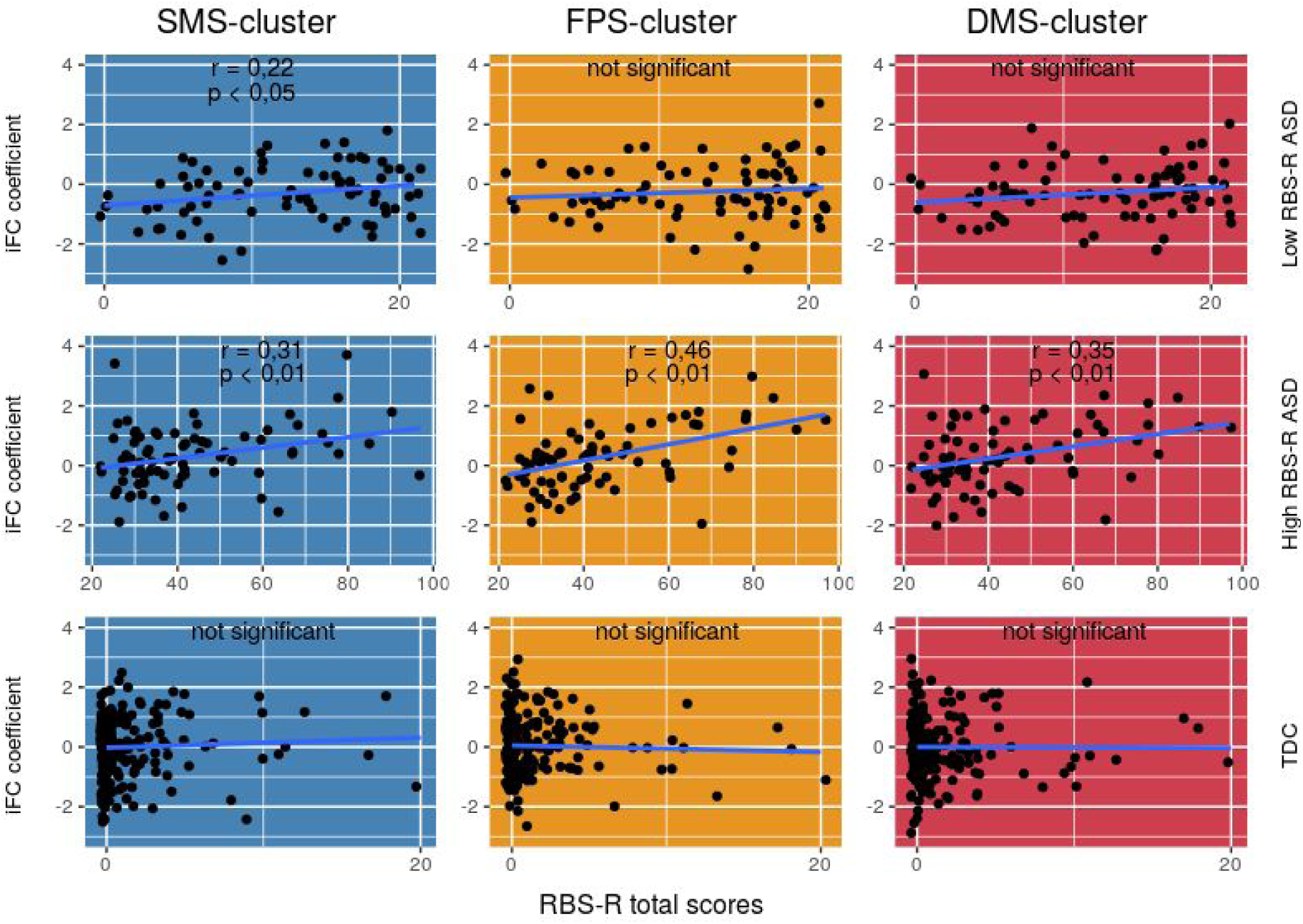
Scatterplots of correlation between residual z-scores of the iFC of each seed-cluster group and RBS-R total scores for the following samples : a) Low RBS-R ASD sample, b) High RBS-R ASD sample and c) TDC sample.

## Discussion

We used resting state functional connectivity to examine the striatal functional connectivity as a function of restricted and repetitive behaviors in a 157 subject sample of 5 to 17 year-old children and adolescents with ASD. Seed based functional connectivity analyses found new evidence of a correlation between striatal connectivity and RBB. The more severe the RRB were, the stronger the connection between striatal seeds and cortical areas.

First, the more severe Total RRB scores were the stronger the connections between the striatal somato-motor seed and an occipital cortical area. Secondly, the more severe Total RRB scores were the stronger the connections between two striatal seeds, the fronto-parietal and default mode striatal seeds, and a frontal cortex area in the left hemisphere. Thirdly, these connections are not found in a sample of comparable neurotypical subjects. Chen et al. found a similar implication of regions. They linked the connectivity of these same areas as a characteristic of ASD subjects, when compared to controls using the ABIDE cohort (45). They found that out of 220 regions, the most informative connectivity was sourced in the somatosensory, default mode, visual, and subcortical regions. All regions we have more specifically linked to RRB. The association between the connectivity of striatal seeds and the cuneus cortex or the superior frontal cortex linked to RRB is a novel finding. But these regions have been independently linked to ASD or RRB.

Several limitations are to be taken into consideration. Are study excluded low functioning children with ASD, not allowing to generalize our finding to the entire population with ASD. Also, we did not apply a global signal regression, which might have maximized the information available (46).

Furthermore, we used striatal seeds based on adult imaging adults when children and ASD subjects have shown a changing brain activity related to age (47). Finally, we excluded the cerebellum from our study. Cerebellum imaging needs specific setting, and several of the ABIDE site did not include the area.

Further research is needed to link RRB and their neural correlates.

## Conclusion

In sum, the current results demonstrate specific striatal functional connectivity related to the severity of restricted and repetitive behaviors in a large sample of children and adolescents with ASD. More precisely, stronger connections with occipital and frontal cortical areas were detected. These stronger connections were not found in neurotypical subjects. Intrinsic functional connectivity of the striatum could serve as a potential biomarker for improved detection of RRB severity.

